# Oxidative stress inducing agents’ copper and alloxan accelerate cell cycle re-entering of somatic plant cells in the presence of suboptimal exogenous auxin

**DOI:** 10.1101/2020.08.22.263079

**Authors:** Taras Pasternak

**Affiliations:** Institute of Plant Biology, Biological Research Centre, Hungarian Academy of Sciences, Temesvári krt. 62., H-6726, Szeged, P.O. Box 521, H-6701, Szeged, Hungary

## Abstract

The physiological status of differentiated somatic plant cells and kinetics of re-entering in cell cycle were investigated in the case of Medicago sativa leaf protoplasts after the application of oxidative stress-inducing agents. Excess copper (30 *μ*M) and alloxan (0.5 mM) accelerated cell cycle re-entry at an exogenous auxin concentration that alone was insufficient to induce cell activation. Application of stress-inducing agents accelerated changes in the nuclei landscape with further faster re-entry in DNA replication and cytokinesis. This acceleration was accompanied by a lower level of reactive oxygen species (ROS) accumulations. At later stages, stress-agents treated cells resemble stem cells in planta with a smaller size, higher cell viability, lower ROS level, and lower activities of major ROS scavenging enzymes. A similar cellular response could be achieved by increasing the exogenous auxin concentration. Based on these experimental results, it is suggested that sub-lethal stress treatments evoke a transient cell state that accelerates cellular reprogramming. We also speculate that this transient cell state serves as an effective mechanism for protection against oxidative stress.

## 1. Introduction

Environmental stresses induced by pollutants or other factors can negatively affect plant growth and development. In majorities of the cases, such adverse conditions alter the metabolism of reactive oxygen species resulted in it over accumulations [1].

Oxidative stress is defined as a significant imbalance between the generation of ROS and cellular antioxidant cascade activities what lead to a steady-state increase in ROS accumulation and potential damage of the cellular structure [2]. ROS scavenging capacities in planta were dependent on cell types, cell status, and communication between cells. Investigating the effect of stress-induced agents on different cell types and mechanisms of protection on cellular levels is required.

In particular, mesophyll protoplasts, alfalfa leaf protoplast-derived cells, provide an excellent experimental system to study plant cell physiology, including stress-response, de novo cell cycle re-activation and developmental transition [3].

ROS is a key factor in plant stress physiology. ROS are involved in the regulation of numerous cellular processes in plant cells and serve as essential components of many signal transduction pathways [4]. One such process is differentiation/re-activation of somatic plant cells [5], [6]. The importance of ROS generation for plant development was confirmed by experiments where inhibition of ROS generation with DPI (diphenyleneiodonium) or scavenging *H*_2_*O*_2_ by DMTU (dimethylthiourea) inhibited cell cycle activation and plant growth [5]. Similar observations were also reported for animal cells: ROS generation was a trigger to a re-programmed somatic cell to pluripotent cells [7]. In our previous works, we found that combination of low concentration of exogenous auxin with external stressors (either iron, nitric oxide, or paraquat) induced the activation of “embryogenic-type” of cell division in alfalfa leaf protoplasts [8] [5] [9]. However, specific effects of these stressors still not clear and require further clarification. Copper (Cu) is an essential micronutrient for all plant species (Marschner 1995). Namely, copper is a key structural and catalytic component of a large number of proteins, a cofactor in many enzymes, transcription factors and protein interaction domains as Cu/Zn SOD, plastocyanin, ATPase, COPT, etc. (for review see [10], [11]). Copper is also involved in regulating cell wall lignification and ascorbic acid metabolism [12]. Recently it has been found that copper plays a role in epigenetic regulation [13]. Optimal Cu concentration in plant cell culture media is in the range of 0.2-0.5 *μ*M ([14] vs [15]). Besides these functions as a nutrient, copper is a crucial redox-active agent in biological systems as an integral component of several electron transport chains as reductive and oxidative in chloroplasts and mitochondria, correspondingly. Copper plays a dual role in redox regulation [16]: as an integral constituent of Cu/Zn–superoxide dismutase (Cu/Zn-SOD) Cu involves in detoxification of superoxide radicals. However, excess copper eventually leads to the generation of a high amount of ROS through the Fenton reaction [17]. This oxidative burst event has been investigated in detail after applications of excess copper (100 *μ*M) to tobacco BY2 cells [18]. Those cells, which are not able to effectively scavenge ROS have permanent oxidative stress. Numerous publications reported the inhibitory effect of copper on cell cycle progression in animal/human and plant cells. In thymus and spleen cells, 25 *μ*M copper increased the size of the sub-population of G0/G1 cells, what the result of inhibition of cell cycle progression [19] was. Similarly, copper caused cell cycle arrest in liver cells [20]. In the root meristem of Allium cepa, copper (8 *μ*M) inhibited the cell cycle progression [21]. In Arabidopsis plants, 25 *μ*M copper prevented root apical meristem development but induced lateral root formation [22]. Interestingly, the effect of excess copper on shoot development was much less pronounced. The last paper suggests a possible involvement of the copper in de novo cell cycle activation. It fits well with [6], who demonstrated that 30 *μ*M copper-induced cell cycle re-entry of stationary phase cultured Medicago cells. Alloxan is another potential inducer of *H*_2_*O*_2_ generation in animal and plant cells [23] [24] [25] [26] [27]. Application of alloxan to Arabidopsis seedlings inhibited the growth of the main root but promoted lateral root induction [25]. However, the direct effect of alloxan on cell cycle in plants has not been investigated yet. Majority of investigations of connections between cell cycle progression and ROS were performed either on cell suspension [28] or on whole plant level [21] when cell cycle progression is dependent from redistribution of phytohormone auxin [29]. Much less attention has been paid on an investigation of the link between cell physiological status, chromatin accessibility, and cell cycle activation during cell reprogramming.

In the current study, we investigated how oxidative stress-inducing agents’ copper and alloxan interact with suboptimal exogenous auxin influencing chromatin accessibility, cell cycle activation/progression, and ROS generation/scavenging in plant cells at different stages of cell cycle re-entering. Our hypothesis is that non-lethal dose of stress agents induced only very transient ROS accumulation, which may trigger physiological processes including ROS scavenging, cell cycle activation, and induction of autonomous development dependent on auxin concentrations.

Our observations revealed that copper/alloxan in combination with exogenous auxin-induced rapid activation of ROS scavenging activity prevents ROS accumulation, and activated cellular re-programming, starting with chromatin relaxation. Stress treatment-induced formation of small, densely cytoplasmic cells with lower vacuolar pH. This cell type has a high developmental potency and is considered to have embryogenic competence [8] Fehér et al. 2003, [9].

The observation that these cells after prolong incubation have lower ROS level and lower antioxidant enzyme activities (including lower level of ascorbate and glutathione) to compare with cells cultured without stress-agents suggests that cells in this developmentally competent transient cell state are protected from oxidative damage due to the prevention of ROS generation/accumulation beside the activity of ROS scavenging.

## 2. Materials and Methods

### 2.1. Protoplasts cultures

Protoplasts were isolated from 4–6-weeks-old alfalfa (Medicago sativa L. ssp. varia; embryogenic genotype A2) plants grown in vitro as described else-where (Pasternak et al. 2000, 2002). Purified protoplasts were washed twice with W5 solution and re-suspended at a cell density of 105 protoplasts/ml in a modified K75 medium supplemented with 0.4 M glucose, 0.1 M mannitol, and the appropriate growth regulators and stress agents. Copper as CuSO4×5H2O and alloxan (A4713 Sigma-Aldrich) at appropriate given concentrations were added before medium sterilization by filtration. Stress treatments were applied from the beginning of protoplast culture. All treatments were carried out at 6 biological replicates for Cu and 5 for alloxan.

### 2.2. Determination of cell cycle kinetics and viability

Cell viability was determined with Evan’s blue [30]. The frequency of cells that already passed through their first cell division was determined by microscopical observations inspecting at least 500 cells per sample. For the determination of the number of cells that passed the DNA replication phase, cells were cultured in the presence of 10 *μ*M of 5-bromo-2^*/*^-deoxyuridine (BrdU; Amersham Biosciences, Vienna, Austria), added at 12 h after starting the culture. Samples were taken at the indicated time points. Immunological detection of BrdU in isolated nuclei was carried out using standard protocols (Amersham Biosciences, Vienna, Austria), as described previously in detail [31] [8]. DNA stainability was detected by flow cytometry. Briefly, nuclei were isolated from the cells fixed in Galbraith buffer [32] containing 2% formaldehyde for 30 minutes. Nuclei were washed twice with buffer, stained with 5 mg/l propidium iodide, and separated on a fluorescence-activated cell sorter (FACS) machine with constant running parameters [9]. The position of the peak in flow cytometry reflected DNA stainability (amount of dye that can bind to open sites of DNA).

### 2.3. Hydrogen peroxide measurement

The determination of the concentration of hydrogen peroxide released from the cells per 1 h was done according to [5]. The rate of cellular *H*_2_*O*_2_ was expressed as *μ*mol *H*_2_*O*_2_ from 10^6^ viable cells per hour.

### 2.4. Protein extraction and enzyme activity determination

For protein isolation, cells were frozen in liquid nitrogen and stored at –70*°*C until analysis. Frozen samples were ground in a minimal volume of ice-cold extraction buffer containing 100 mM sodium phosphate (pH 7.0), 5 mM ascorbate (only in the case of APX), 1 mM EDTA, 1 mM PMSF, 2 mM 2-mercaptoethanol and 1 mM leupeptin. Extracts were centrifuged for 15 min. at 14,000 g at 4*°*C. The supernatant was collected and protein concentration was determined using the Bio-Rad protein assay kit (BIORAD, Hercules, USA) and bovine serum albumin as a standard. Ascorbate peroxidase (APX; EC 1.11.1.11) activity was determined spectrophotometrically by monitoring the decrease in absorbance at 290 nm in 1 ml reaction mixture, containing 50 mM potassium phosphate buffer (pH 7.0), 0.5 mM ascorbate, 0.1 mM *H*_2_*O*_2_, 0.1 mM EDTA and 10 *μ*g protein [33]. A calibration curve was made measuring the absorbance of the reaction mixture (without extract) with 5–200 *μ*M ascorbate. Catalase (CAT; EC 1.11.1.6) activity was determined by monitoring the decomposition of *H*_2_*O*_2_ at 240 nm in 1 ml reaction mixture, containing 66 mM potassium phosphate buffer (pH 7.0), 12.5 mM *H*_2_*O*_2_ and 10 *μ*g protein [34] and defined as OD240/*μ*g protein/min. The enzyme activities measured right after protoplast isolation were highly variable from experiment to experiment likely due to slight differences in conditions of the explants and isolation procedures. That is why the relative changes in enzyme activities were calculated and are shown using the initial values (after protoplasts isolation) as reference 1.

### 2.5. Vacuolar pH determination

The pH-sensitive fluorescent dye FDA (Sigma) was dissolved in dimethyl sulfoxide at the concentrations of 2 mM. For dye loading, small aliquots of the cells (approximately 200 *μ*L of culture) were transferred to 1.5-mL reaction tubes, and the appropriate amount of the dye to final concentration 2 *μ*M was added. Cells were gently transferred to microscopic slides with a jacket and cover with a coverslip. Fluorescence was measured after 10 min after loading. Fluorescent signals were detected by a fluorescent microscope (Axiovert 135 M; Zeiss). For ratiometric pH measurements, the excitation was at 440 and 490 nm. Fluorescence passing through a 515-nm dichroic mirror and 540 nm bandpass filter was recorded with a CCD camera. A 25% transmittance neutral density filter was used between the light source and the filter to decrease excitation energy and minimize photo-bleaching. Excitation time was kept to 0.125 s to minimize dye bleaching. In this case, photobleaching of the dyes represented less than 5% of fluorescence per observation/scan. Autofluorescence represented less than 1% of the total signal from dye-loaded protoplasts and did not change with time or experimental treatment. Background fluorescence intensity (together with dark camera level) was measured based on the average background signal from each individual image from an area next to the samples and was subtracted from the fluorescent intensity of the cells. The arithmetic operation was used to distinguish the pH-dependent fluorescence obtained at 470 nm excitation from the pH-independent fluorescence obtained at 440 nm excitation.

### 2.6. Data analysis and statistics

All experiments were carried out in five independent biological repetitions. Statistical analyses of data were performed by either T-tests (comparison single points) or one way ANOVA using Microsoft Excel software.

## 3. Results

### 3.1. Effect of protoplasts isolation procedure on cell physiology

First, it was investigated how the procedure of protoplast isolation affected the physiological status of the mesophyll cells. We have found that incubation of the leaf segments in the enzyme solution and all washing steps led to a significant reduction in antioxidant enzyme activities (APX and POD) (Fig. 1). This, in turn, led to the rise of *H*_2_*O*_2_ level in the cells immediately after isolation.

**Figure 1:**
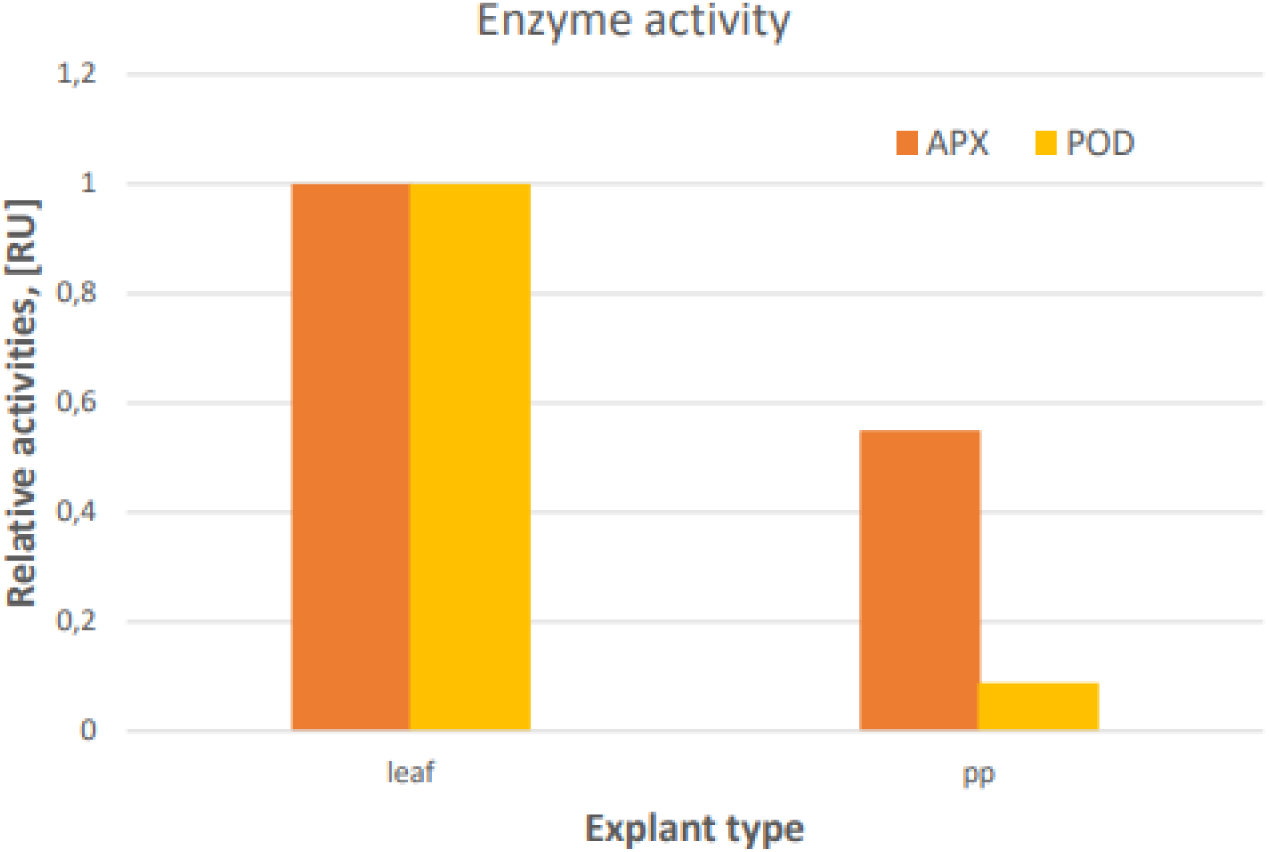
Effect of protoplasts isolation procedure on activity of antioxidant enzymes.

### 3.2. Copper application accelerates cell cycle re-activation in an auxin-dependent manner

Copper (30 *μ*M) has been applied to freshly isolated Medicago leaf protoplasts in combination with two different concentrations of the auxin (2,4-D): 0.05 mg/l (appr. 220 nM) and 0.2 mg/l (appr. 880 nM). In both treatments, extra copper had a significant effect on several cellular features, including DNA stainability and cell cycle progression (Fig. 2). Copper, combined with exogenous auxin, increased DNA stainability (Fig. 2a), which means the amount of propidium iodide which can bind to DNA, indicating that the chromatin was in a more accessible status (Pasternak et al., 2000). The positive effect of the copper on DNA stainability was followed by more rapid re-entering in DNA replication that was detected as the ratio of the cells incorporating BrDU into their DNA during a given incubation period (Fig. 2b). Correspondingly, the number of cells, which passed cytokinesis at 96 hours was also higher in the presence of copper (Fig. 2c).

**Figure 2:**
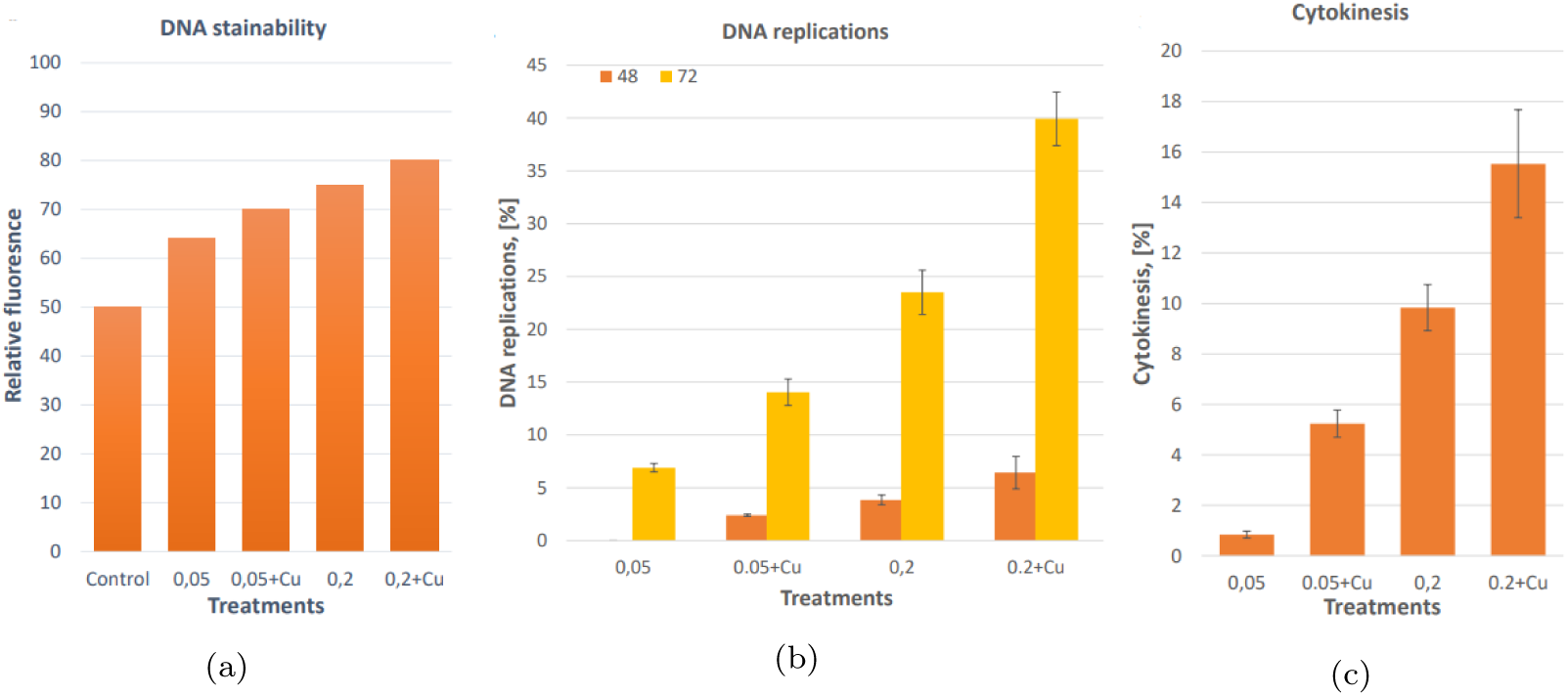
Effect of the copper on cell cycle reactivation. Medicago leaf protoplasts were cultured on the medium contain 0,05 and 0,2 mg/l (0,25 *μ*M and 1 *μ*M) 2,4-D with and without 30 *μ*M Cu added as CuSO_4_. (a) - position of the G0/G1 peak on the nuclei histogram. Nuclei were isolated from cells after corresponding treatments. Nuclei were stained with propidium iodide and subjected to run on flow cytometer. Position of the peak corresponding to 2C DNA contents were determined (Y-axis). Control is nuclei from freshly isolated protoplasts; (b) - percent of the cells what pass DNA replications at given time point. BrDU was added at 12 hours after culture start, cells were fixed at given time points and BrDU was detected with specific antibody. Number of BrDU positive cells were divided to total number of cells in the population; (c) - percent of the cells what pass cytokinesis at 96 hours after isolation.

Next, it was asked whether the application of copper resulted in oxidative stress. To clarify this point, we measured the *H*_2_*O*_2_ level, the activities of key antioxidant enzymes, and cell viability. It was found that the *H*_2_*O*_2_ level was significantly lower in the case of copper application (Fig. 3a) at all time points likely because of higher APX activities as detected after 12 and 36 hours of cultivation Fig. 3c. However, after 96 hours of cultivation in the presence of copper, higher APX/CAT activities were detected only under 0.05 mg/l 2,4-D, but not under 0.2 mg/l 2,4-D.

**Figure 3:**
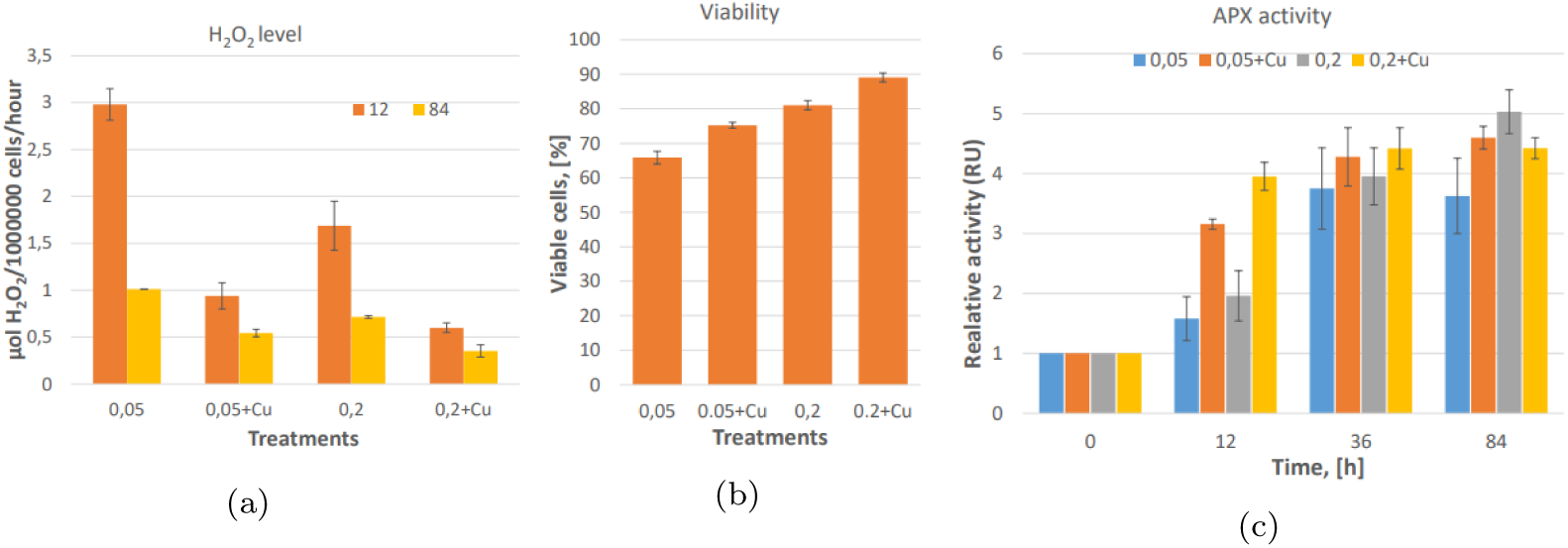
Effect of Cu on ROS accumulation, cell viability and APX activities. (a) - amount of non-scavenging *H*_2_*O*_2_ realizing to the culture medium; (b) - cell viability determined by Evans Blue; (c) - total APX activities.

Cell viability was detected as the uptake of Evans Blue correlated with the *H*_2_*O*_2_ level and was significantly higher in the case of copper applications (Fig. 3). Interestingly, under 2 mg/l 2,4-D viability was similar as in the case of 0.2 mg/l 2,4-D supplemented with 30 *μ*M Cu.

Copper also had a significant impact on cell morphology in an auxin-dependent manner -it resulted in the formation of small, densely cytoplasmic cells with small storage vacuoles and compact nuclei. (Fig. 4a, Fig. A.9). These features characterize the cells which grow in the presence of 8.8 *μ*M 2,4,-D and exhibit embryogenic potency [8].

**Figure 4:**
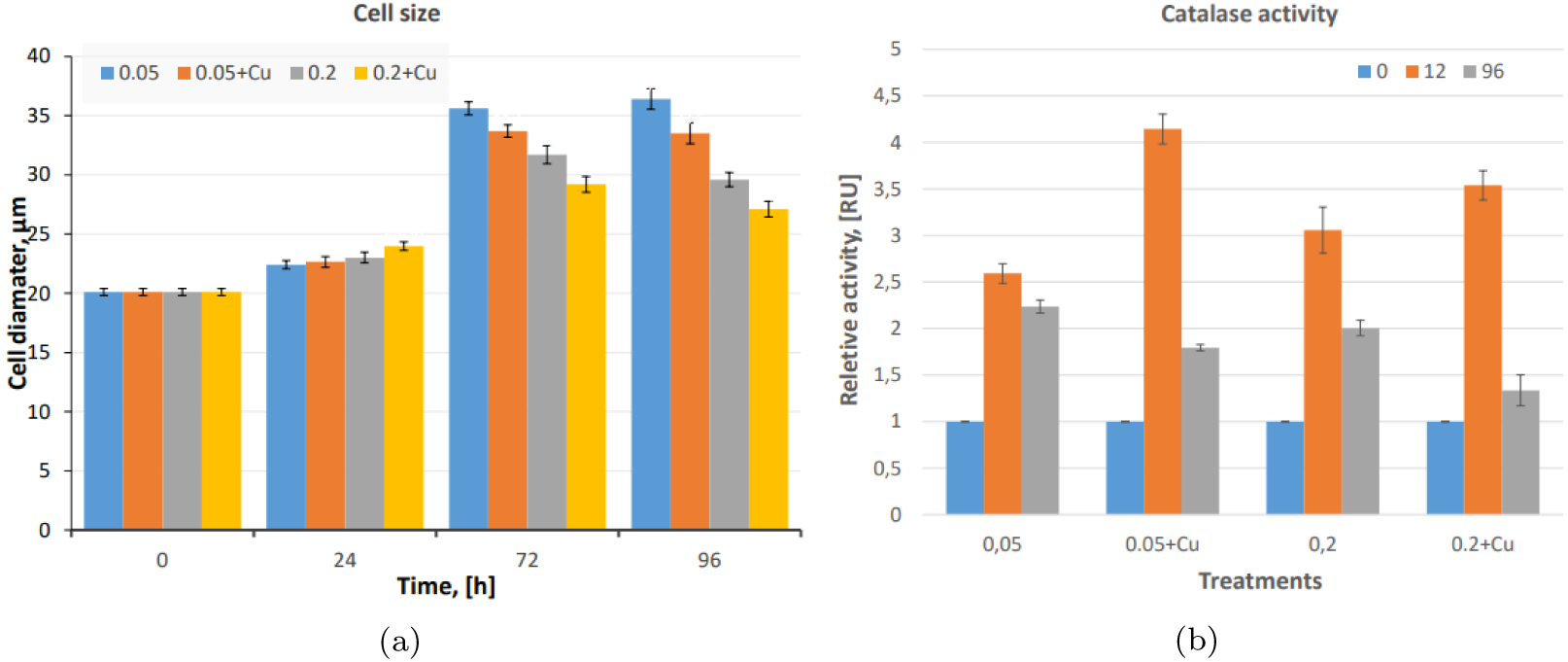
Effect of copper on cell expansion and catalase activity. (a) - cell size; (b) - catalase activity. Activity shown in relative units with activity in fresh protoplasts as 1 unit.

### 3.3. Alloxan promotes cell cycle re-entering and the formation of compact cells

Since copper can be considering as an essential nutrient, we used another stress-induced agent - alloxan - to prove the general role of stress application in cell cycle reactivation.

As it is shown on Fig. 5, application of 0.5 mM alloxan had similar effects on cell activation and morphology as 30 *μ*M copper: it accelerated chromatin relaxation as indicated by the volume of nuclei and nucleoli (Fig. 5a Fig. 5b), increased the ratio of the cells passing DNA replication and cytokinesis (Fig. 5c Fig. 5d). Higher cell viability (**??**), what was observed after 96 hours of cultivation, was accompanied by lower *H*_2_*O*_2_ accumulation (Fig. 6b) and lower activities of the key ROS-scavenging enzymes (APX, Fig. 6c), POD, catalase. Significantly smaller cells were formed (Fig. 6d, Fig. A.8) characterized by lower vacuolar pH (Fig. A.9).

**Figure 5:**
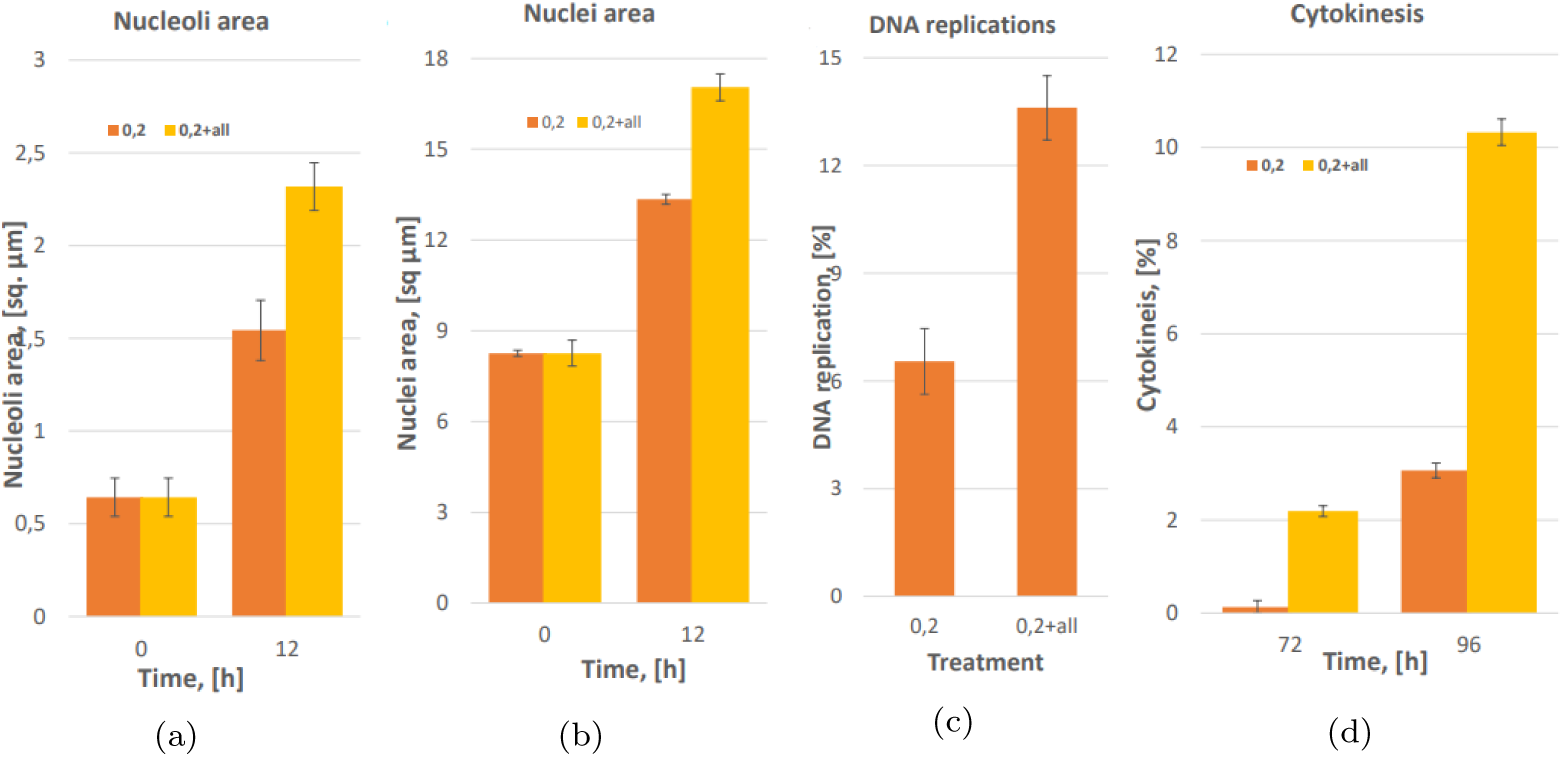
Effect of alloxan on nuclei morphology and cell cycle progression. (a) - nucleoli area; (b) - nuclei area; (c) - percent of the cells what pass DNA replications after 60 h; (d) - percent of the cells what pass cytokinesis at 72 and 96 hours after isolation.

**Figure 6:**
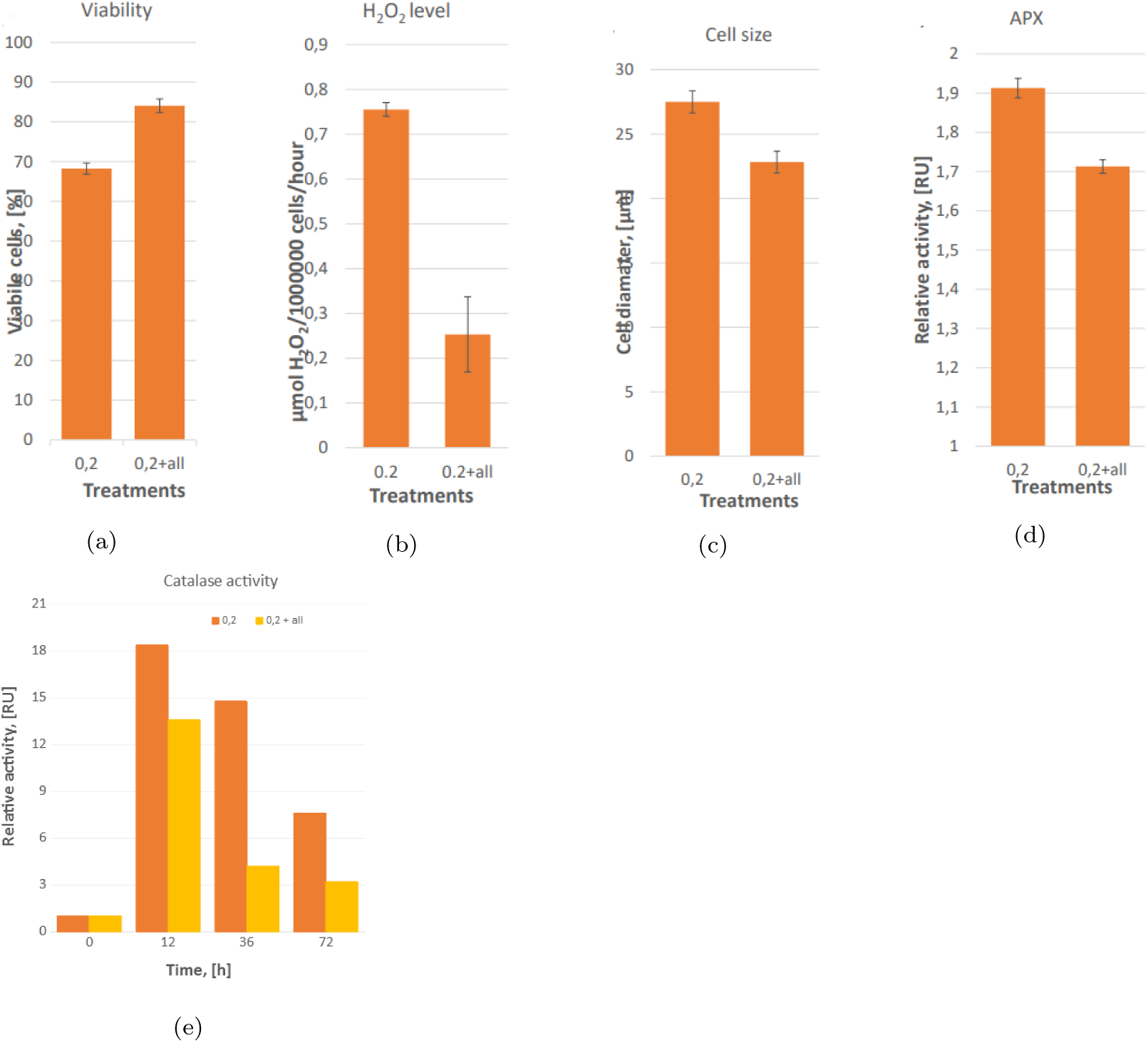
Effect of alloxan on viability, *H*_2_*O*_2_ level and antioxidant enzymes activities. (a) - cell viability after 96 h; (b) - *H*_2_*O*_2_ level after 96h; (c) - cell size after 96h; (d) - APX activity after 96 h; (e) - catalase activity kinetics during cultivation

Altogether, one can conclude that there was a general similarity between the two stress-inducing agents concerning their effect on protoplasts growing in the presence of suboptimal auxin concentration.

### 3.4. Stress pre-treated cells require suboptimal auxin concentration for further microcolony growth

Since cells after stress-treatments have features of stem-like cells in planta, which are semi-autonomous ea. do not require endogenous hormones, we checked the possibility of whether cultured stress after stress-treatment became relatively independent from exogenous auxin. After 96 hours in the culture, cells were washed from stress-induced agents embedded in alginate and cultured in the presence of very low auxin concentration. It was shown that cell after alloxan/Cu treatment were able to develop further as a compact globular colony, while cell from the control cannot (Fig. A.10).

## 4. Discussion

### 4.1. Stress-inducing agents can promote cell cycle re-entering

Excess copper can be considered a stress-inducing agent, which may lead to slow-down cell cycle progression rate as frequently observed in stressed plant cells (for review: [35]). In accordance, in both plant and animal cells, stress was reported to inhibit cell cycle progression [28] [36]. However, [22] demonstrated that copper, while inhibited cell cycle progression in the main root apical meristem, induced de novo lateral root induction (de novo cell cycle activation). Similarly, alloxan in Arabidopsis inhibited main root growth but induced lateral root induction [25]. This raised the possibility that the application of stress-induced agents has a different effect on cell cycle progression and cell cycle induction, respectively. Copper was shown to increase the activity of the cell-cycle-regulatory cyclin-dependent-kinase (CDK) complex during the re-activation of stationary phase cell cultures [6]. In the copper moss Scopelophila cataractae copper mediates auxin signaling to control cell differentiation [37].

Physical access to DNA is a highly dynamic property of chromatin that plays an essential role in establishing and maintaining cell activity [38]. In differentiated somatic cells, cell activity is very restricted mainly to photo-synthesis. That’s why the conversion of these cells to cell cycle active requires significant chromatin re-organization. The ability of DNA assessment can be quantified by measure the amount of dye that can be binding to nuclear DNA. Here we demonstrated that two stress-inducing agents (copper and alloxan) in the presence of the external auxin were able to accelerate chromatin relaxation. This was reflected by higher DNA stainability and a rapid increase in nuclei/nucleoli size. Interestingly this effect mimics that of higher auxin concentration. It fits well with the finding that stress can increase internal auxin concentration in the cells [8]. This statement also fits well with the recent finding that copper increased auxin contents in maize [39]. We asked the next question: does excess copper and alloxan-induced oxidative stress (accumulation of excess reactive oxygen species)? In order to determine the level of oxidative stress, the rate of excess *H*_2_*O*_2_ and the activities of the main enzymatic *H*_2_*O*_2_ scavengers, ascorbate peroxidase, and catalase (for review [40]) were determined in protoplasts-derived cells at different time-points after stress-agent applications. The amount of *H*_2_*O*_2_ released into the medium, spatially and temporally separated from the sites of its production, reflects the amount of *H*_2_*O*_2_, which could not be scavenged by the antioxidant system (ROS-scavenging enzymes and low molecular weight antioxidants) in the cells [5]. We demonstrated that treatment with copper and alloxan does not induce the higher accumulation of *H*_2_*O*_2_ to compare with corresponding control because of higher ROS-scavenging activities. It means that stress-agents induced acceleration of changes in chromatin structure (size and stainability of nuclei) (Fig. 2a, Fig. 5a, Fig. 5b) and cell cycle progression (DNA-replication and mitosis) accompanied by less *H*_2_*O*_2_ accumulation to compare with the corresponding control cells.

The lower level of *H*_2_*O*_2_ may be related either to higher scavenging or lower production rate. Therefore, we analyzed the activity of main scavenging enzyme activities and have found that the lower *H*_2_*O*_2_ accumulation level in stress-agents treated cells correlated well with higher APX, POD and CAT activities after 12 and 36 hours. Interestingly, after 96 hours of lower *H*_2_*O*_2_ level in stress-agents treated cells in combination with 0.05 mg/l 2,4-D correlated with higher ROS scavenging system activities, while in combination with 0.2 mg/l 2,4-D with lower scavenging capacity. This phenomenon can be explained by the lower level of ROS generation in stem-like cell types. Stem-like cells in planta were characterized by small size because of small protein storage vacuole. In our hand, both copper and alloxan treatment led to the formation of similar vacuole type with higher vacuolar pH as in the non-treated cells. This phenomenon linked well with the finding of [41], who showed that copper negatively regulated V-ATPase activity and reduced vacuolar pH and vacuolar size in yeast. Next, we analyzed the sites of ROS generation/scavenging. There are two main ROS scavenging enzymes in plant cells: catalase (CAT), abundant in peroxisomes, and several isoforms of APX (ascorbate peroxidase) localize in chloroplasts and the cytoplasm. We found that after an initial break of the activity of both enzymes during protoplast isolation, there was a switch from chloroplastic to cytoplasmic APX (and general downregulation of CAT activity. The switch from CAT-to APX-activity has been described previously in protoplast-derived cells of grape [42]. Our finding also confirmed the results of [43] that *H*_2_*O*_2_ level declined in tobacco protoplasts after cell re-programming but remained high in the grapevine protoplasts, which did not re-program. Type of cell division on the medium with 0.2 mg/l 2,4-D plus copper/alloxan exhibited features of stem-like meristematic cells in planta: exceedingly small size, small storage vacuoles, and no functional chloroplasts. In contrast, cells which were not exposed to stress factors had large size, bigger not-fused chloroplasts, and a large vacuole and can resemble non-stem meristem cells in planta. Cells in SAM/buds in planta can initiate new organs, and they serve as auxin source through internal auxin synthesis. Given the cell similarity between stem cells in planta and cells after copper/alloxan treatments, we ask questions about the hormonal-independence of these cells and the possibility of further morphogenesis. Our results demonstrated that short-term stress application induced morphogenic potential of the cells, which were able to further proliferate to compact globular colony (Fig. A.10).

The above results demonstrated that stress-inducing agents copper and alloxan (in combination with suboptimal auxin concentration) accelerated chromatin relaxation and, in this way, accelerated cell cycle progression. Moreover, the application of stress-inducing agents did not induce ROS accumulation accompanied by rapid activation of the ROS scavenging system or, at the later stage, by the reduction in ROS generation. The second option can be considered as a non-canonical way of protection against stress by inducing relatively dormant stem-like cells that in planta can be resembled by dormant bud. On the contrary, cells that do not get auxin or have only low auxin can possess a canonical way of protection by scavenging ROS.

## 5. Conclusions

Our study demonstrated that:

1. Application of stress-induced agents in combination with suboptimal auxin concentration accelerated chromatin relaxation in mesophyll protoplast-derived cells;
2. Application of stress-inducing agents in non-lethal dose in combination with the amount of external auxin that is suboptimal to support cell division led to the transient accumulation of ROS what was diminished by higher activity of ROS scavenging enzymes.
3. Stress-induced agents combined with higher auxin concentration led to the formation of stem-like cells that demonstrated low ROS levels with low levels of antioxidant machinery that may suggest the existence of the non-canonical way of protection against stress agents as a transition to stem-like cells.

## Appendix A. Supplemental Figures

**Figure A.7:**
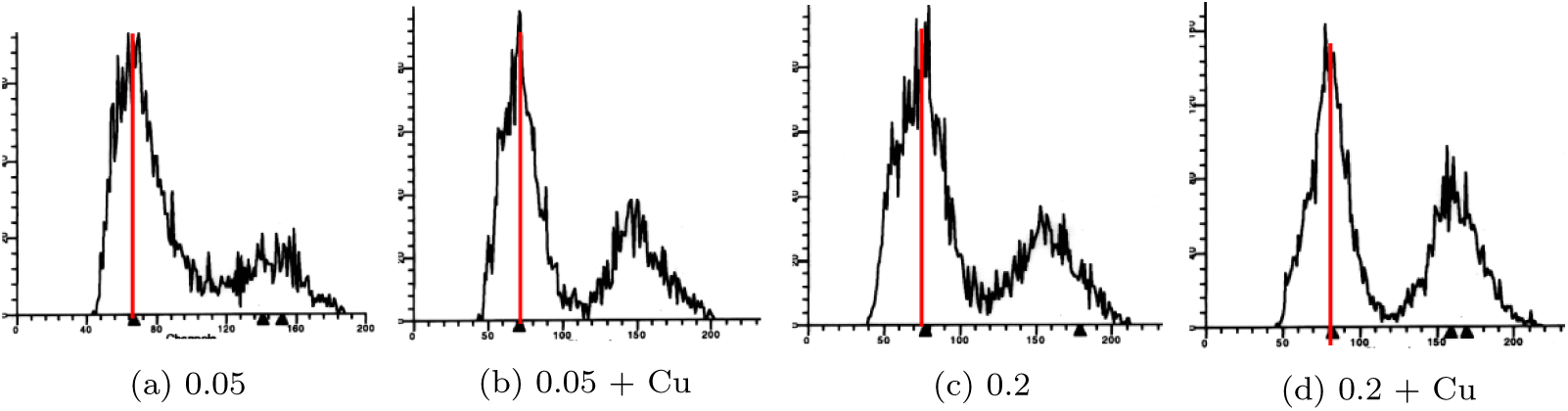
Flow cytometry histogram after 96 hours in culture.

**Figure A.8:**
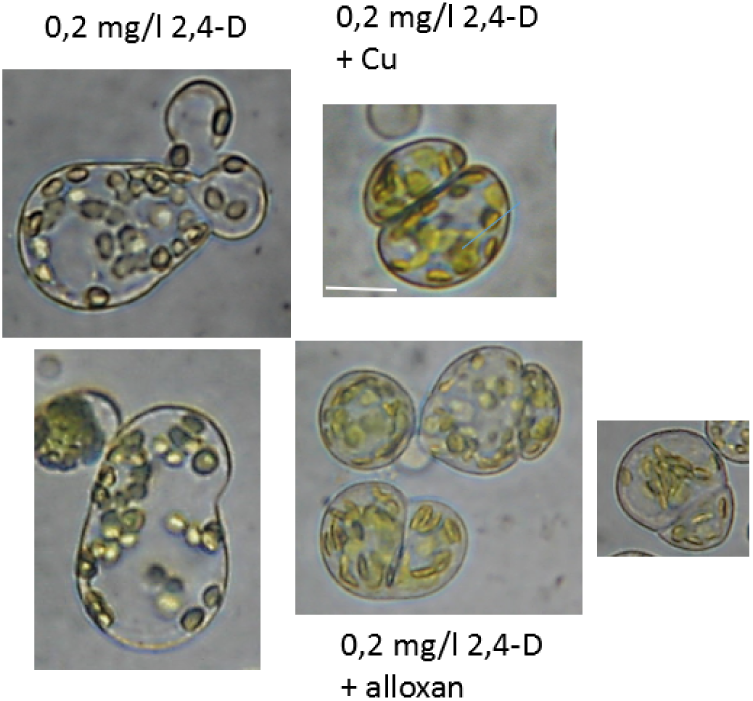
Cells morphology after 96 hours in culture. Scale bar- 10 *μ*m.

**Figure A.9:**
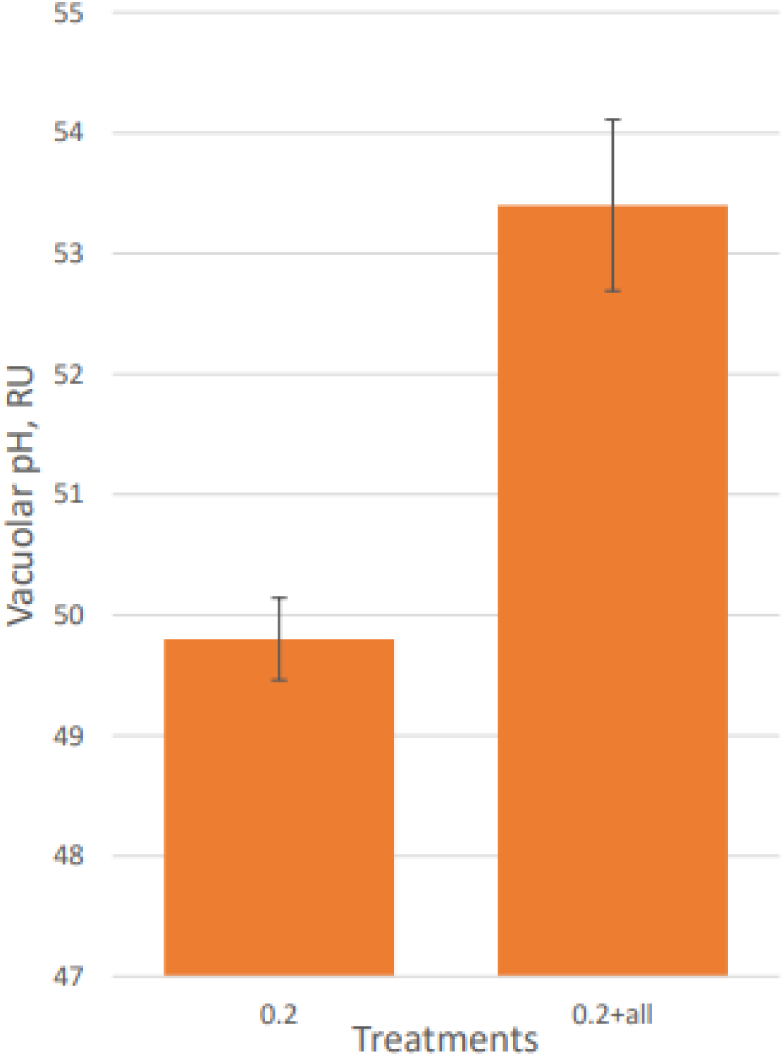
Vacuolar pH

**Figure A.10:**
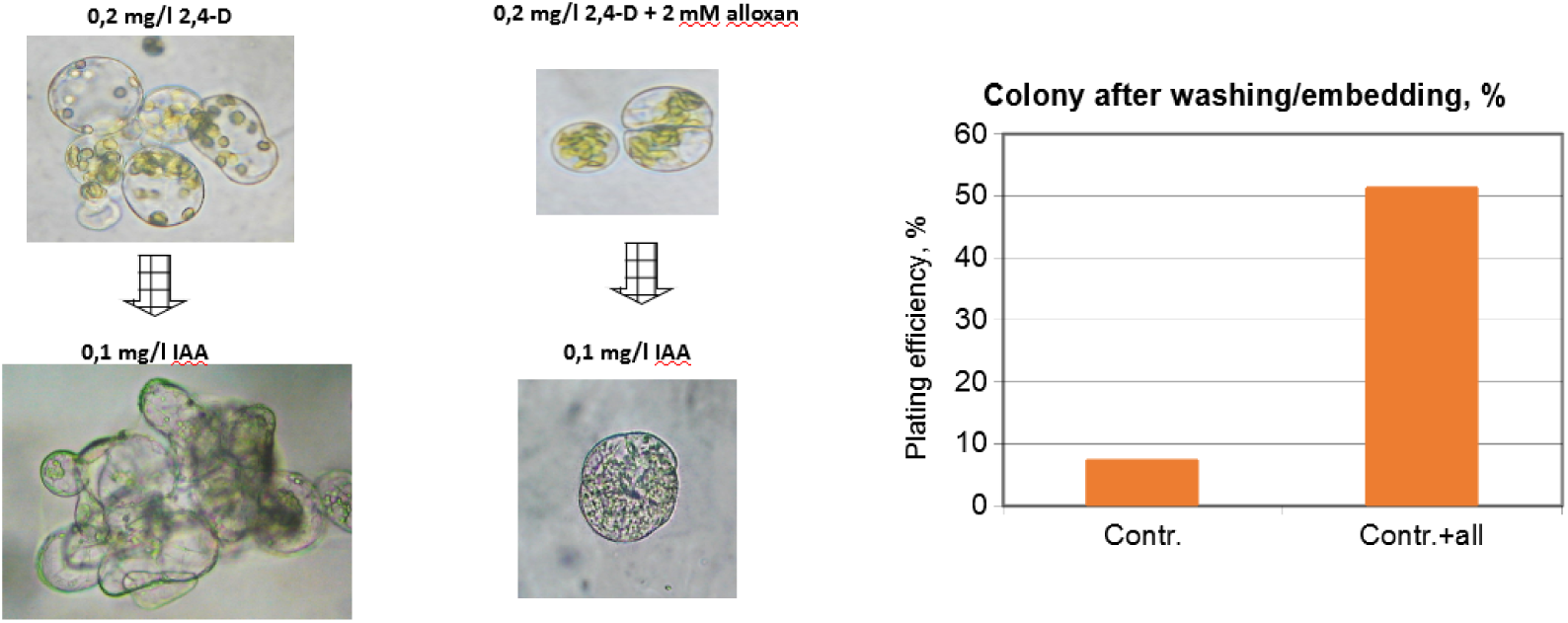
Auxin-independency of protoplasts-derived colony.

